# The role of hippocampal CAMKII in resilience to trauma-related psychopathology

**DOI:** 10.1101/2022.07.05.495828

**Authors:** S. Hazra, J. D. Hazra, R. Amit Bar-On, Y. Duan, S. Edut, X Cao, G Richter-Levin

## Abstract

Traumatic stress exposure can form persistent trauma-related memories. However, only a minority of individuals develop post-traumatic stress disorder (PTSD) symptoms upon exposure. We employed a rat model of PTSD, which enables differentiating between exposed-affected and exposed-unaffected individuals. Two weeks after the end of exposure, animals were tested behaviorally, following an exposure to a trauma reminder, identifying them as trauma ‘affected’ or ‘unaffected’. In light of the established role of hippocampal synaptic plasticity in stress and the essential role of Ca2+/calmodulin-dependent protein kinase II (CaMKII) in hippocampal based synaptic plasticity, in two separate experiments, we pharmacologically inhibited CaMKII or knocked-down αCaMKII in the dorsal dentate gyrus of the hippocampus (dDG) following exposure to the same trauma paradigm. Both manipulations brought down the prevalence of ‘affected’ individuals in the trauma-exposed population. A day after the last behavioral test, long-term potentiation (LTP) was examined in the dDG as a measure of synaptic plasticity. Trauma exposure reduced the ability to induce LTP, whereas, contrary to expectation, αCaMKII-kd reversed this effect. Further examination revealed that reducing αCaMKII expression, enables the formation of αCaMKII-independent LTP, which may enable increased resilience in the face of a traumatic experience. The current findings further emphasize the pivotal role dDG has in stress resilience.

## 1. INTRODUCTION

PTSD is recognized as a major health challenge, but there are only partially effective treatments for the disorder. Often, PTSD co-occurs with mood and anxiety-related disorders and is associated with a severe disability and medical illness (1, 2). One significant obstacle hindering clinical solutions to PTSD resides in the individual variability that exists in susceptibility to trauma, which leads some to develop PTSD while others seem to be resilient even in the case of severe traumatic experiences (3, 4). Epidemiological data demonstrated that even after experiencing severe, traumatic stress, only about 7% of the exposed individuals would display symptoms of post-traumatic stress disorder (PTSD) (5, 6). When exploring risk factors that may explain this individual variability, childhood adversity in humans (7) and juvenile stress in animal models were found to exacerbate the impact of adult stress on activity and anxiety-like behaviors (8–10). Elucidating the neuronal mechanisms responsible for trauma susceptibility or resilience will unravel key molecules or pathways that can be targeted to promote resilience.

PTSD has been repeatedly documented to involve a reduced hippocampal functioning (14) and volume (16, 17). Experimental evidence indicates that the dorsal hippocampus, which corresponds to the human posterior hippocampus, is specifically involved in the formation of stable ‘declarative’ memory (18) and in encoding cognitive and spatial information (19–21). The ventral or temporal pole of the hippocampus, which corresponds to the anterior hippocampus in humans, modulates emotional and affective processes such as stress responses and fear behavior (21–23). Nevertheless, pharmacological and genetic manipulation derived studies have demonstrated a role for the dorsal hippocampus, and particularly the dorsal dentate gyrus (dDG), in mediating stress effects on hippocampal functioning (24). Altered expression of GABAergic factors (25–27) and altered local circuit activity within the dDG (28, 29), have been implicated in response to stress and in stress resilience. In particular, the hippocampal DG was shown to respond differently to various stressors, enabling it to integrate both emotional and cognitive aspects of an experience and share that data with the hippocampus proper, as well as with a network of brain regions such as the amygdala and prefrontal cortex (8, 24).

Long-term potentiation (LTP) is suggested to serve memory formation (30, 31). Stress is well known to modulate hippocampal LTP as well as hippocampus-dependent learning and memory. Alterations of plasticity in the hippocampus, amygdala, and medial prefrontal cortex were found to play a pivotal role in stress vulnerability, pathology, and stress resilience (3, 32). Exposure to stress differently alters neural plasticity in these regions. For example, while exposure to stress was constantly shown to impair LTP in the CA1 field of the hippocampus, the plasticity aberrations induced by stress in the DG are much less consistent (33–36).

One leading player in the induction and maintenance of LTP in the hippocampus is Ca^2+^/calmodulin-dependent protein kinase II (CaMKII) (39–41). The mammalian CaMKII is a dodecameric holoenzyme containing 12 subunits. Four genes encode CaMKII- α, β, γ, and δ that exhibit alternative splicing in their variable linker domain. The α and β isoforms are predominant in the hippocampus (42, 43). Because of its prominent role in mechanisms of neural plasticity and the potential role of aberrant neural plasticity under stressful conditions, several studies have examined the potential role of the αCaMKII in anxiety-related behaviors. Overall, the findings indicate that manipulating the expression or functioning of αCaMKII affects anxiety symptoms, though there are still many inconsistencies that need further investigation (46–50).

Considering the impaired memory, learning and neural plasticity in the hippocampus of PTSD subjects (51), and the significant role CaMKII plays in neuroplasticity, we hypothesize that αCaMKII may induce a pathological plasticity in the hippocampus resulting in distortion of memories and maladaptive enhancement of emotional events. The main goal of this study is to examine whether alterations in CaMKII expression will influence the percentage of PTSD affected individuals. Specifically we focus on reducing αCaMKII expression in the dDG as this sub-region of the hippocampus has been implicated in defining stress vulnerability or resilience (8,24,27,52,53). Furthermore, in order to relate to individual variability in response to trauma (4), we employed our recently developed behavioral profiling approach (54–57), which enables dissociating trauma-exposed-affected from trauma exposed- unaffected individuals.

## 2. RESULTS

### 2.1. Combined juvenile stress and UWT (JS-UWT) increases the prevalence of ‘affected’ animals

We have earlier shown that pre-exposure to JS exacerbates the effects of exposure to trauma in adulthood, increasing the percentage of the affected individuals (8,10,55). Thus, we applied a PTSD paradigm consisting of JS and UWT in adulthood (Experiment 1, Supporting information). The behavioral responses of individual animals were tested in the OF and EPM following re-exposure to the UWT associated odor cue (55). Looking first at the group averages of the behavioral tests, revealed that JS-UWT animals showed a significant reduction in the total distance covered in the OF (t=3.75, p<0.001) and in the EPM (t=4.46, p<0.001), compared to that of the CTR group (Fig S1 B, C). Additionally, there was an enhanced anxiety-like behavior indicated by the reduction in the anxiety index both in the OF (t=6.90, p<0.001) and the EPM (t=4.73, p<0.001), measured as the percentage of distance covered in center divided by total distance (OF) and percentage of distance covered in the open arms divided by the total arms distance (EPM).

Since some individuals are more susceptible to develop an affective symptomatology after a traumatic experience, a comparison of the different groups’ means would not reveal differences at an individual level between subjects that coped with trauma and those who developed pathology. Hence, we applied the behavioral profiling approach (55, 56) to identify vulnerable and resilient individuals during the exploration tests. Behavior profiling analysis, which employs a dedicated algorithm, including fifteen behavioral parameters extracted from the OF and the EPM tests (see “Methods” in the Supporting information for a detailed description), identified affected and unaffected individuals. Comparisons between the groups using Pearson χ2 analysis revealed that the proportion of affected rats in the JS-UWT group was significantly higher (83.0%) compared to that of the control group (5.0%), (X2 = 123.5, p<0.001) (Fig S1D).

### 2.2. Reducing CaMKII functionality in the dDG of trauma exposed animals decreased the prevalence of ‘affected’ individuals

Next, we examined whether alterations in CaMKII functionality specifically in the dDG would affect the percentage of affected individuals. For this purpose, we implemented two different intervention strategies: pharmacological inhibition of CaMKII by KN93, and a local viral-vector-mediated knockdown of αCaMKII expression (Experiments 2 and 3).

#### 2.2.1. Inhibition of dDG CaMKII by KN93 reduced the prevalence of “affected” individuals

One-way ANOVA revealed a significant effect for groups in total distance traveled in the OF (F (2, 49) = 5.627, p= 0.006) and EPM (F(2, 49) = 10.564, p< 0.001), as well as in the anxiety index measured in the OF (F(2, 49) = 9.099, p< 0.001) and EPM (F(2, 49) = 40.83, p< 0.001; Fig 1 B-C). As expected, exposure to JS-UWT increased anxiety levels both in the OF and EPM, as denoted by the reduction in the total distance covered in the OF and EPM compared to that of the saline-injected, non-trauma-exposed group (Post-hoc Bonferroni p= 0.017 and p< 0.001, respectively). A similar effect was found for the anxiety index in both behavioral tests (Post-hoc Bonferroni p< 0.001 for OF and EPM). The anxious-like behavior in the OF and EPM was counteracted by the injection of the CaMKII inhibitor KN-93 prior to subjecting the rats to the adulthood trauma. KN-93 significantly increased the total distance covered in the OF vs. the saline-injected JS-UWT group (Post-hoc Bonferroni p= 0.019), and a positive trend was observed in the relative time spent in the center of the arena. The results in the EPM were compatible with those in the OF, both for total distance and relative time spent in the open arms (Post-hoc Bonferroni p= 0.0013 and p < 0.001, respectively), suggesting that local inhibition of CaMKII in the dDG of adult rats is enough to attenuate their anxiety-like behavior. In line with the above, behavioral profiling analysis revealed a significantly higher proportion of affected rats in the JS+UWT group (76.0%) compared to that of the control group (X2 = 98.507, p <0.001;). KN93 to the dDG before UWT, significantly increased resilience of the JS-UWT exposed rats, raising the proportion of unaffected individuals from 24.0% to 91.0% (X2= 69.31, p<0.001) compared to JS-UWT-exposed animals injected with saline (Fig 1D).

**Figure 1.**
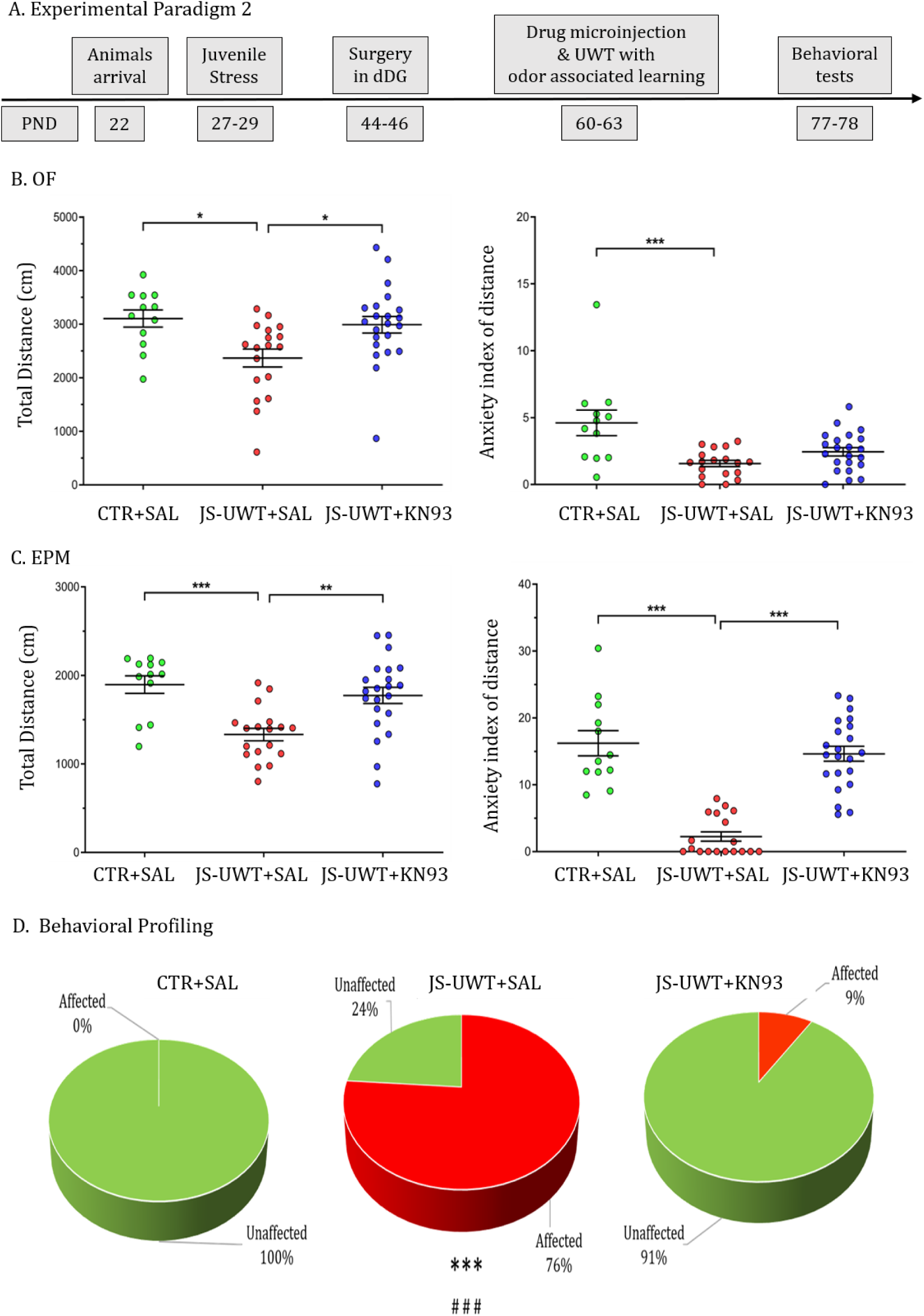
Administration of CaMKII inhibitor KN93 in the dDG reduces the prevalence of affected animals after exposure to juvenile stress and UWT as adulthood stress (JS-UWT). See supporting information for further details. (A) Schematic representation of experimental timeline, (PND-postnatal day; supporting information). (B) and (C) Averaged group effects in the open field and the elevated plus maze respectively. (Left) Total distance travelled in the maze and (Right) Anxiety index of distance. All values are mean ± SEM ***p<0.001, ** p<0.01, *p<0.05. (D) Behavior profiling analysis shows a significant higher proportion of affected animals among JS- UWT+SAL rats compared to that of the CTR+SAL population, whereas the JS- UWT+KN93 group shows significant reversal effect. Values are the % affected and unaffected animals in each group. ***p<0.001, ###p<0.001.

#### 2.2.2. The impact of knockdown of αCaMKII in the dDG

##### 2.2.2.1 Verification of the efficacy of viral knockdown of CaMKII expression in vitro and in vivo

To verify the efficacy and specificity of the viral vector we assessed viral transfection efficiency and αCaMKII knockdown efficacy *in vitro* (Fig 2A-C). Three shRNA sequences aimed at reducing αCaMKII expression were examined in 293 cell lines. The three constructs show similar transfection efficacy as determined by EGFP fluorescence. Subsequent western analysis revealed robust reduction of αCaMKII expression compared to that of the control cells, which overexpressed a construct carrying a sequence for αCaMKII without any of the shRNA carrying vectors, and results showed that shRNA1 had the best knockdown effect. We further tested the efficacy of the shRNA αCaMKII AAV vector *in vivo* within dDG. Only animals with correct injection sites on both hemispheres were included in the analysis, as ascertained by GFP expression (Fig 2D). Local injection of shRNA αCaMKII AAV into the dDG resulted in 56% knockdown of endogenous αCaMKII expression (t=3.606, df=13, p=0.003) (Fig 2E).

**Figure 2.**
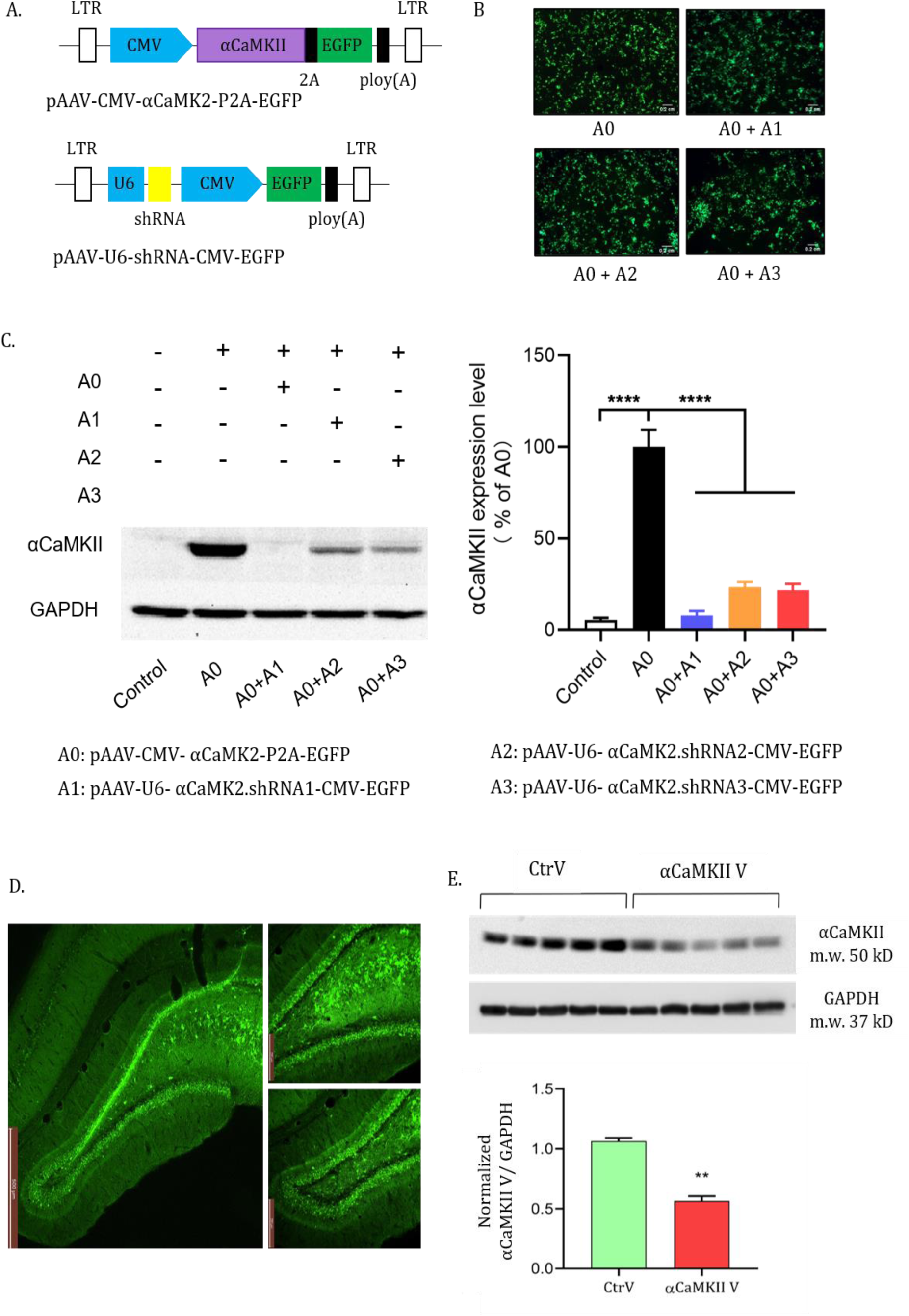
Validation of αCaMKII virus. The knockdown efficiency of three shRNAs in 293 cell line. (A) up: Schematics of AAV construct for overexpressing (pAAV-CMV-αCaMK2-P2A-EGFP); down: Schematics of AAV construct for knockdown αCaMKII (pAAV-U6-shRNA-CMV-EGFP). ITR, inverted terminal repeats; CMV, cytomegalovirus promoter. (B) Representative fluorescence images of 293 cells after AAV vectors transfection. EGFP, green. (C) αCaMKII protein level in 293 cells transfected with pAAV-CMV-αCaMK2-P2A-EGFP and pAAV-U6-shRNA-CMV-EGFP vectors. (D) αCaMKII virus expression in the dDG. (E) Representative western blot images showed significant reduction in αCaMKII knockdown group. All values are mean ± SEM, ***p<0.001, ** p<0.01, *p<0.05.

##### 2.2.2.2 knockdown of αCaMKII in the dDG reduces the prevalence of “affected” individuals

Animals were injected with αCaMKII shRNA-AAV vector into the dDG two weeks after JS (PND 44-46). They were later subjected to UWT in adulthood (PND 63) and tested in the OF and EPM two weeks later (PND 77-78). In the OF, no group effect was evident in the arena path length, although an apparent effect in total distance was seen in the EPM following exposure to JS-UWT (One-way ANOVA F (2, 44) =7.746 p=0.001). In line with the previous findings, a significant effect for groups was found for the anxiety index measured both in the OF (F (2, 44) =9.303, P<0.001) and EPM (One-way ANOVA (F(2, 44) = 12.50, p< 0.001) (Fig 3 B, C). As expected, exposure to trauma increased anxiety level as marked by the reduction in the total distance covered in the EPM (Post-hoc Bonferroni p= 0.001) and anxiety index in both OF and EPM (Post-hoc

**Figure 3.**
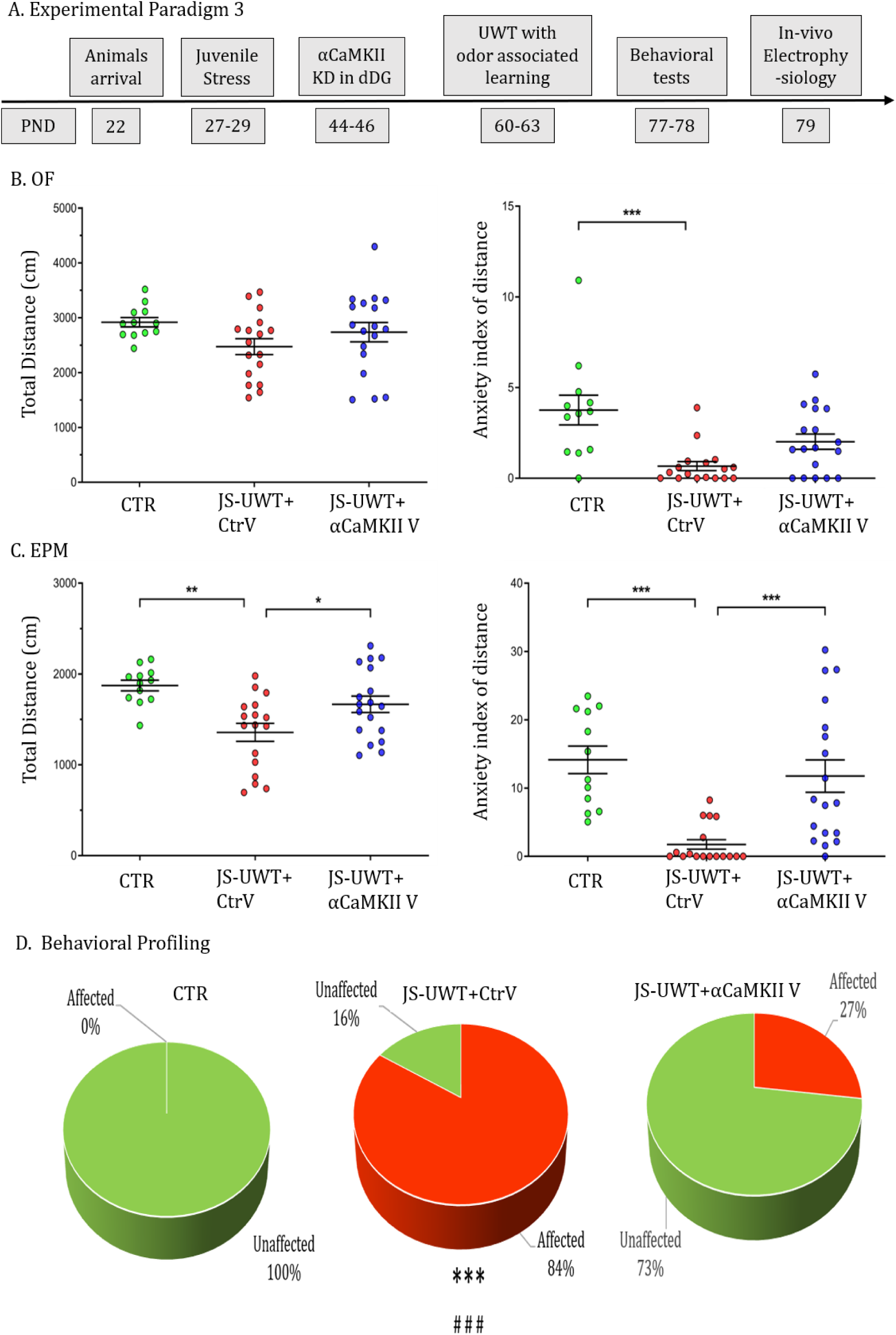
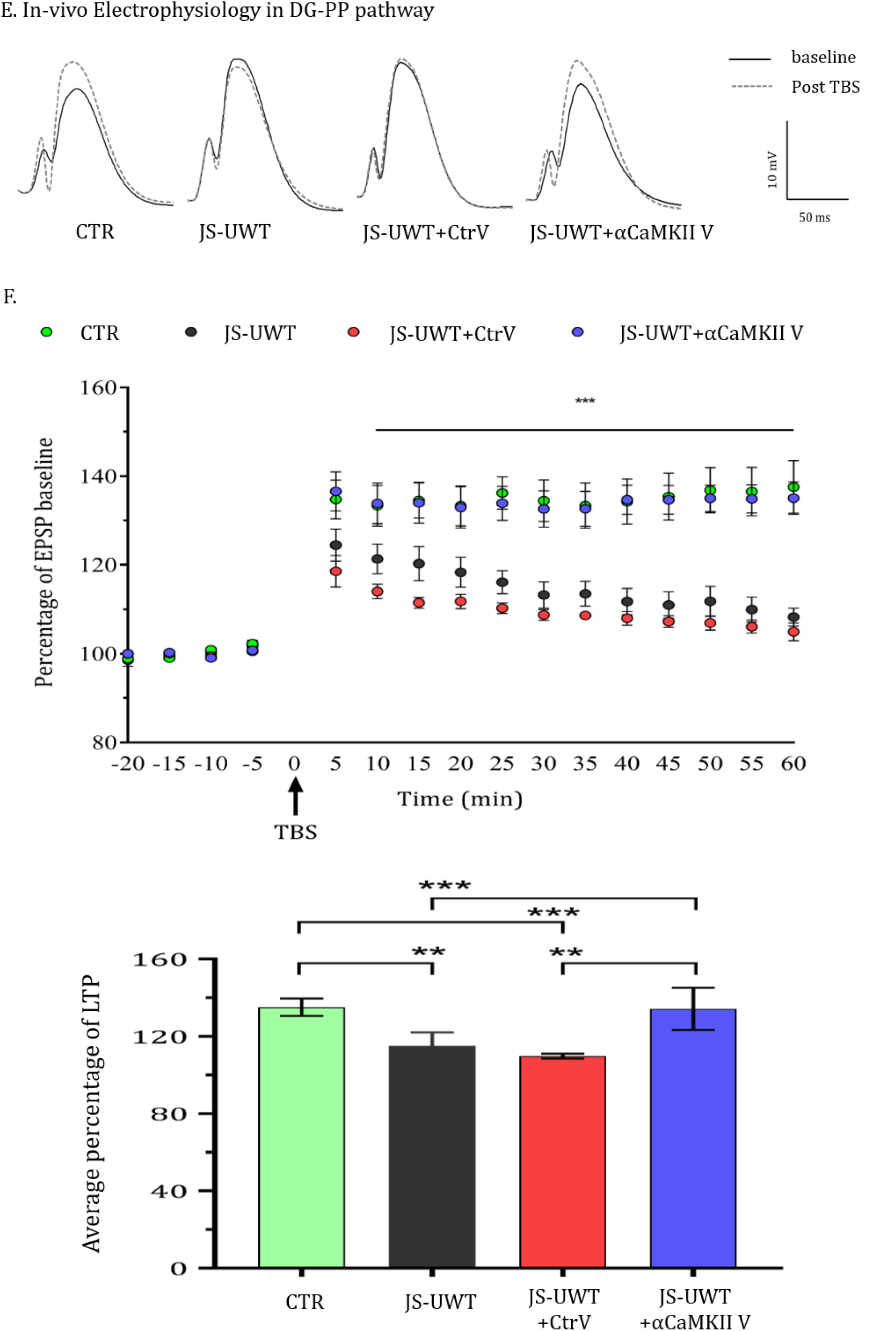
Knock-down of αCaMKII in the dDG reduces the prevalence of affected animals after exposure to JS-UWT and suppressed LTP. See supporting information for details. (A) Schematic representation of experimental timeline, (PND-postnatal day; supporting information). (B) and (C) Averaged group effects in the open field and the elevated plus maze respectively. (Left) Total distance travelled in the maze and (Right) Anxiety index of distance. All values are mean ± SEM, ***p<0.001, ** p<0.01, *p<0.05. (D) Behavior profile approach shows a significant higher proportion of affected animals among JS-UWT+CtrV rats compared to the CTR population, whereas in JS-UWT+ αCaMKII V group the affected population is significantly less compared to JS-UWT+CtrV group. Values are the % affected and unaffected animals in each group. ***p<0.001, ###p<0.001. (E) Effect of αCaMKII knockdown on synaptic plasticity in the DG of the hippocampus. Evoked field potential response recorded in the DG of CTR (n=8), JS-UWT (n=8), JS-UWT+CtrV (n=10), and JS-UWT+αCaMKII V (n=10) (F) A significant DG field potentiation was recorded after TBS in CTR group and JS-UWT+ αCaMKII V, while TBS failed to induce potentiation in JS-UWT and JS-UWT+CtrV group (left), changes in average percentage of LTP after TBS (right). LTP was assessed upon theta-burst stimulation (TBS) to the perforant path-DG pathway, and recording in the dDG. No significant difference in baseline recording was found prior to TBS. Average percentage of LTP (right), The results are the Mean ± SEM. ***p<0.001, ** p<0.01.

Bonferroni P<0.001 and p<0.001, respectively), compared to the control group. Post-hoc test did not find any significant changes after the knockdown of αCaMKII in total distance (p=0.61) and anxiety index (p= 0.12) in the OF, whereas in the EPM, a prominent significant effect for total distance and anxiety index was observed), compared to that of JS- UWT group injected with a control virus (Post-hoc Bonferroni P=0.042 and p=0.001, respectively). In line with the above, behavioral profiling analysis revealed a significantly higher proportion of affected rats in the JS -UWT group (84.0%) compared to that of the control group (F; X2 = 144.828, p <0.001). Targeted knockdown of αCaMKII in the dDG significantly increased resilience of the JS-UWT exposed rats, raising the proportion of unaffected individuals from 16.0% to 73.0% (X2= 65.776, p<0.001) compared to JS-UWT-exposed animals injected with control virus (Fig 3D).

### 2.3. Selective αCaMKII Knock-down enabled LTP in dDG in exposed animals

Twenty-four hours after the last behavioral test, LTP was examined in the dDG in all four groups as a measure of synaptic plasticity. There was no significant difference in the input-output curve between groups, indicating no baseline difference in excitability (data not shown). No significant difference in baseline recording was found prior to TBS. Robust theta-burst stimulation (TBS) was applied to the perforant path (PP)-DG pathway and its impact was recorded in the dDG. After TBS, mixed model repeated measures of ANOVA showed a significant difference between groups (group effect F (3, 28) = 15.44, p < 0.001; time effect; F (2.159, 60.442) = 7.22, p< 0.001; time X group; F (6.476, 60.442) = 3.44, p< 0.001). JS-UWT reduced the ability to induce LTP (Fig 3 E, F). Selective knockdown of αCaMKII reversed this effect in the JS-UWT, and potentiation level was similar to control. Post-hoc comparisons between the four groups revealed that JS-UWT and JS-UWT+Ctr V groups were significantly different from CTR (Bonferroni, p= 0.002 and p< 0.001 respectively) and from JS-UWT+αCaMKII V (Bonferroni, p= 0.002 and p< 0.001 respectively). No significant difference was found between CTR and JS-UWT+αCaMKII V group.

### 2.4. αCaMKII knockdown inhibits LTP induced by regular TBS protocol but not by robust TBS protocol

In the findings above there was a correlation between the behavioral results and the LTP results, such that JS-UWT led to pathological symptoms, and to impaired LTP, while knocking down dDG αCaMKII reduced pathological symptoms and enabled normal levels of LTP. While this correlation appears to suggest a link between the two, the LTP results are not in agreement with previous reports suggesting that hippocampal LTP is dependent on αCaMKII activation (58–60). Thus, recovery of the ability to induce LTP by reducing expression of αCaMKII requires consideration. We set out to examine the possibility that the ability to induce LTP in αCaMKII KD animals was because we applied a robust TBS protocol. For that, we compared the effect of αCaMKII KD on LTP induced by either a robust or a regular TBS protocol, in animals that were not exposed to any behavioral protocol. The results support our assumption showing that αCaMKII V impaired LTP following the regular TBS protocol. No potentiation was evident after 60 min, and levels did not differ from those of TBS baseline. There was no significant difference in the input-output curve between groups, and baseline recording prior to TBS (data not shown) (Fig 4B, C). Repeated measure of ANOVA shows a significant group effect F (2, 22) = 28.09 p<0.001; time effect F (1.802, 39.66) = 4.37, p<0.001; but no time x group effect F (3.605, 39.66) = 2.06, p>0.05). Post-hoc comparisons revealed that the αCaMKII V group was significantly different from the CTR and Ctr V groups (p< 0.001). Moreover, the comparison of averaged potentiation across time points after induction also showed significant difference (one-way ANOVA, F (2, 22) = 21.09 p<0.001, post-hoc Bonferroni CT vs αCaMKII V (p< 0.001 and Ctr V vs αCaMKII V, p< 0.001). In contrast, LTP induced by robust TBS was maintained in all three groups (CTR, n=6; Ctr V, n=6; and αCaMKII V, n=7) (Fig 4D, E). Repeated measure of ANOVA revealed no significant group effect F (2, 16) = 2.530 p>0.05; time effect F (1.996, 27.94) = 1.146, p>0.05; time x group effect F (3.991, 27.94) = 0.687, p>0.05). Furthermore, the comparison of averaged potentiation across time points after induction also showed no significant difference between groups.

**Figure 4.**
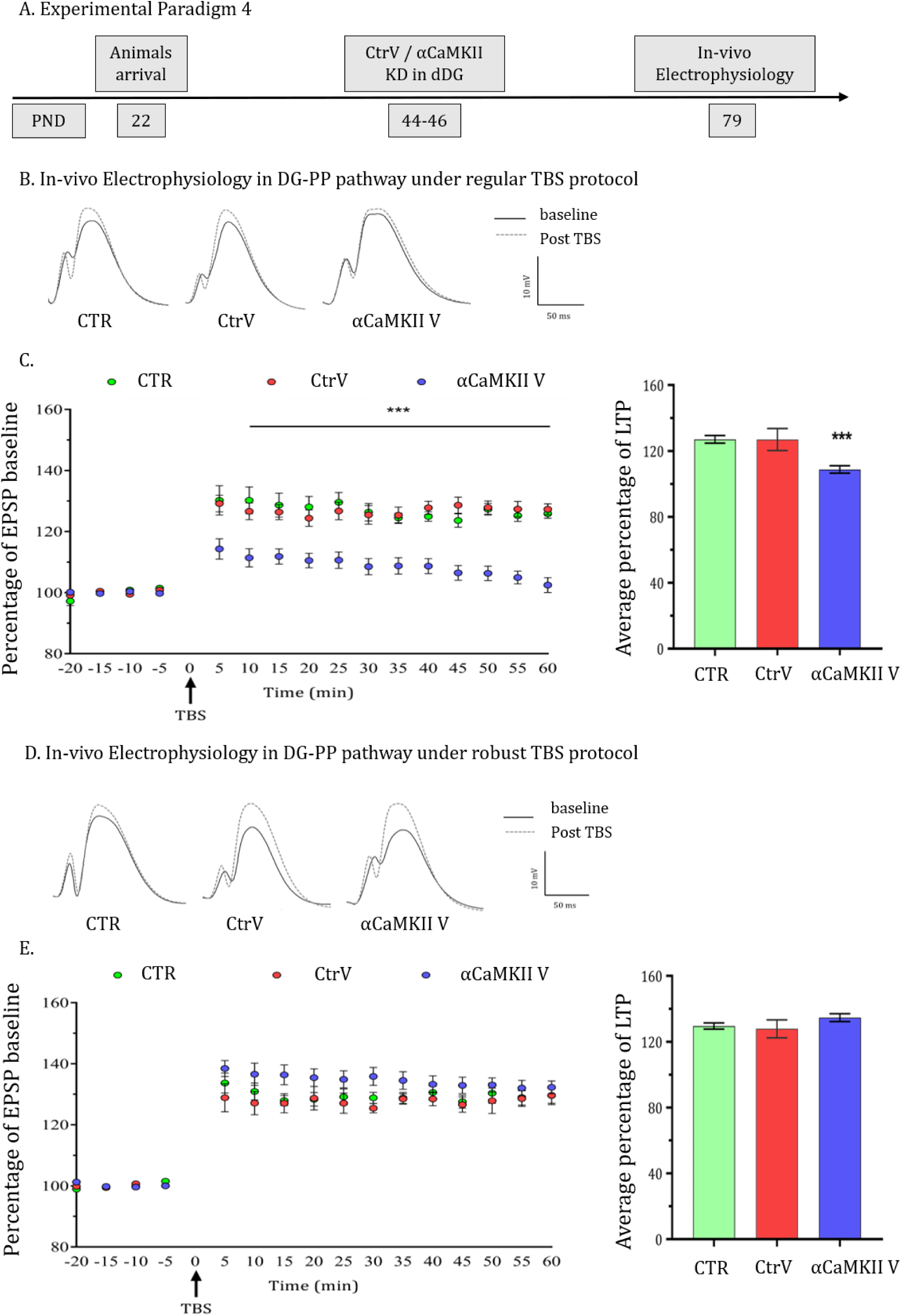
LTP activity under regular or robust TBS protocol following viral inhibition of αCaMKII. (A) Schematic representation of experimental timeline, (PND-postnatal day; supporting information). (B) Evoked field potential response recorded in the dDG of CTR (n=7), JS-UWT+CtrV (n=7), and JS-UWT+αCaMKII V (n=8) under regular TBS protocol. (C) A significant DG field potentiation was recorded after TBS in CTR and CtrV groups, while in αCaMKII V group, TBS failed to induce potentiation (left); changes in average percentage of LTP after TBS (right). (D) Evoked field potential response recorded in the DG under robust TBS protocol, CTR (n=7), JS-UWT+CtrV (n=7), and JS-UWT+αCaMKII V (n=8). (E) TBS induced a significant potentiation of DG field potentials in all the groups (left); no significant changes were found in average percentage of LTP after TBS (right). LTP was assessed upon TBS to the perforant path-DG pathway, and recording in the dDG. No significant difference in baseline recording was found prior to TBS. The results are the Mean ± SEM, ***p ≤ 0.001.

## 3. DISCUSSION

PTSD has been repeatedly documented to involve reduced hippocampal functioning and decreased hippocampal volume (61), with a suggested emphasis on the DG (24, 62). Studies focusing on the dorsal hippocampus have found an impact of stress in that region, particularly relating to resilience, expression of GABAergic factors and local circuit activity in the DG (27, 53).

The involvement of CAMKII in hippocampal plasticity and the role of pathological hippocampal plasticity in PTSD raises the possibility for a role of CAMKII in stress and trauma, but the nature of its involvement is not clear. In a αCaMKII-GFP transgenic mouse line, the αCaMKII-GFP was found to be mostly expressed in the DG region and CA1/CA3. About 70% of granule and pyramidal neurons strongly expressed GFP in the hippocampus (63). According to previous studies, transgenic upregulation of αCaMKII in the forebrain could cause aggression and anxiety-like behavior in mice, indicating a correlation between αCaMKII level and the state of emotion (49). Overexpression of αCaMKII in the medial prefrontal cortex and DG impaired behavioral flexibility and NMDAR-dependent long-term depression, respectively (64, 65). Other studies also reported that overexpression of active form of CaMKII in CA1 of the hippocampus may impair context discrimination- a feature also related to PTSD (66). A single immobilization stress was found to increase phospho-CaMKII levels in the hippocampus, though there was no change in hippocampal levels of CaMKII (67). However, following a single prolonged stress exposure, which is considered an animal model of PTSD, the histone deacetylase inhibitor, vorinostat, ameliorated impaired fear extinction, and this effect was mediated in part by increased expression of hippocampal CaMKII (50). In the current study we aimed to focus on the role of dDG αCaMKII in PTSD, because the dDG was implicated in defining stress vulnerability or resilience (8,24,27,52,55,68).

The lifetime prevalence of exposure to severely stressful events like combat, accidents, natural disasters, assault, or rape is as high as 75–80%, yet only about 10–20% of the population exposed to these stressors will eventually develop PTSD (5). As in humans, not all animals develop PTSD-like symptoms following trauma exposure. The search for neural mechanisms associated with the pathology or with resilience requires separating between those individuals who were affected and those who did not develop pathology following trauma exposure (4). Towards that end the behavior profiling analysis approach was developed. Separating affected and unaffected animals was found pivotal, enabling the identification of individual responses to trauma which would have otherwise been masked by the analysis of averaged group responses (53,55–57). Building on previous findings, we have now developed an advanced, algorithm-based behavioral profiling analysis (see details in Supporting information). The analysis of the behavioral outcome of exposure to JS-UWT corroborates previous findings, demonstrating an increased percentage of affected individuals following the combined exposure to JS, as a risk factor and to trauma in adulthood (8).

We employed here two complementary manipulations in order to examine the role of dDG αCaMKII in trauma vulnerability or resilience: local pharmacological inhibition of CaMKII, before UWT exposure, and reduction of αCaMKII expression by a viral vector. Both approaches significantly compromise the potential contribution of dDG αCaMKII to trauma responses. Indeed, the similar outcome resulting from both manipulations greatly increases the confidence in the main finding, indicating that reducing dDG αCaMKII functionality enhances trauma resilience. CaMKII is a highly abundant brain protein concentrated in the postsynaptic density (45, 70), and it is essential in the adult rodent hippocampus for LTP generation (71). We observed impaired plasticity in the DG-PP pathway in affected JS-UWT exposed animals. Prior data suggests that stress can impair cellular plasticity (72, 73), and under some conditions, suppress plasticity in the dDG (28). Diminished plasticity in the DG-PP pathway may reflect altered hippocampal response as observed in PTSD patients (16, 74). Reducing the expression of αCaMKII in JS-UWT exposed animals, prevented the impact of the trauma at both the behavioral and electrophysiological levels, indicating that reduced expression of DG αCaMKII enabled LTP and a form of trauma resilience.

While these results demonstrate an elegant correlation between the behavioral and electrophysiological findings, they also raise an important question: if CaMKII has an essential role in synaptic plasticity, and inhibiting it should block LTP (60,75–77), how come αCaMKII knockdown can lead to recovery of trauma-induced suppression of DG LTP? One possible explanation could be that the use of a strong protocol to induce LTP, as we employed here, enabled the induction of a αCaMKII-independent form of LTP. In order to verify that the result is an outcome of the protocol employed, we tested in a separate group of animals the ability of αCaMKII knockdown to impair LTP when employing a regular protocol. Indeed, LTP was significantly impaired employing a regular TBS protocol, which effectively leads to LTP in control animals (Fig 4C). This result indicates two important points: First, the viral vector used here is effective and functional. Second, as expected, reducing αCaMKII expression impairs the ability to induce LTP in the dDG. However, in another set of animals, reducing the expression of αCaMKII could not block the induction of LTP when applying a strong TBS protocol, a result that is in agreement with the notion that robust TBS enables the induction of a αCaMKII-independent form of LTP.

Accumulating evidence indicates that there are different forms of LTP, with distinct associated mechanisms (78–82). αCAMKII-dependent LTP may be the dominant form of LTP in the hippocampus (83), but other forms of LTP may be induced even when αCaMKII is blocked, or when its expression is inhibited. For example, selective application of CaMKII inhibitors (KN62 or KN93) did not block LTP in rat DG slices. Nevertheless, when KN62 or KN93 were applied together with the PKA inhibitor (KT5720) or the MEK inhibitor (Rp-8-cAMP), DG-LTP was completely blocked, indicating the existence of both αCAMKII-dependent, and αCAMKII- independent forms of LTP (84).

We suggest that under normal conditions, the activation of αCAMKII has a dual effect, supporting the formation of αCAMKII-dependent LTP, but also inhibiting other forms of LTP (Fig 5A). As a result, TBS leads mainly to the formation of αCAMKII- dependent LTP. Exposure to trauma tends to suppress both αCAMKII-dependent LTP and αCAMKII-independent LTP (Fig5B), with an overall result of suppressing LTP. The inhibition of the expression of αCAMKII reduces the probability of activating αCAMKII-dependent LTP, while at the same time it may also remove the αCAMKII- induced inhibition of other forms of LTP. In turn, this inhibitory removal increases the likelihood of triggering these other forms of LTP, despite the inhibitory effect of trauma (Fig 5C). The results obtained by exposure to trauma and reduction in the expression of αCAMKII, are in agreement with the notion that under normal conditions, LTP is mainly a αCAMKII-dependent LTP. They may also provide an explanation for how exposure to trauma may suppress all forms of LTP, while reducing the expression of αCAMKII may enable the formation of αCAMKII-independent forms of LTP in trauma-exposed individuals.

**Figure 5.**
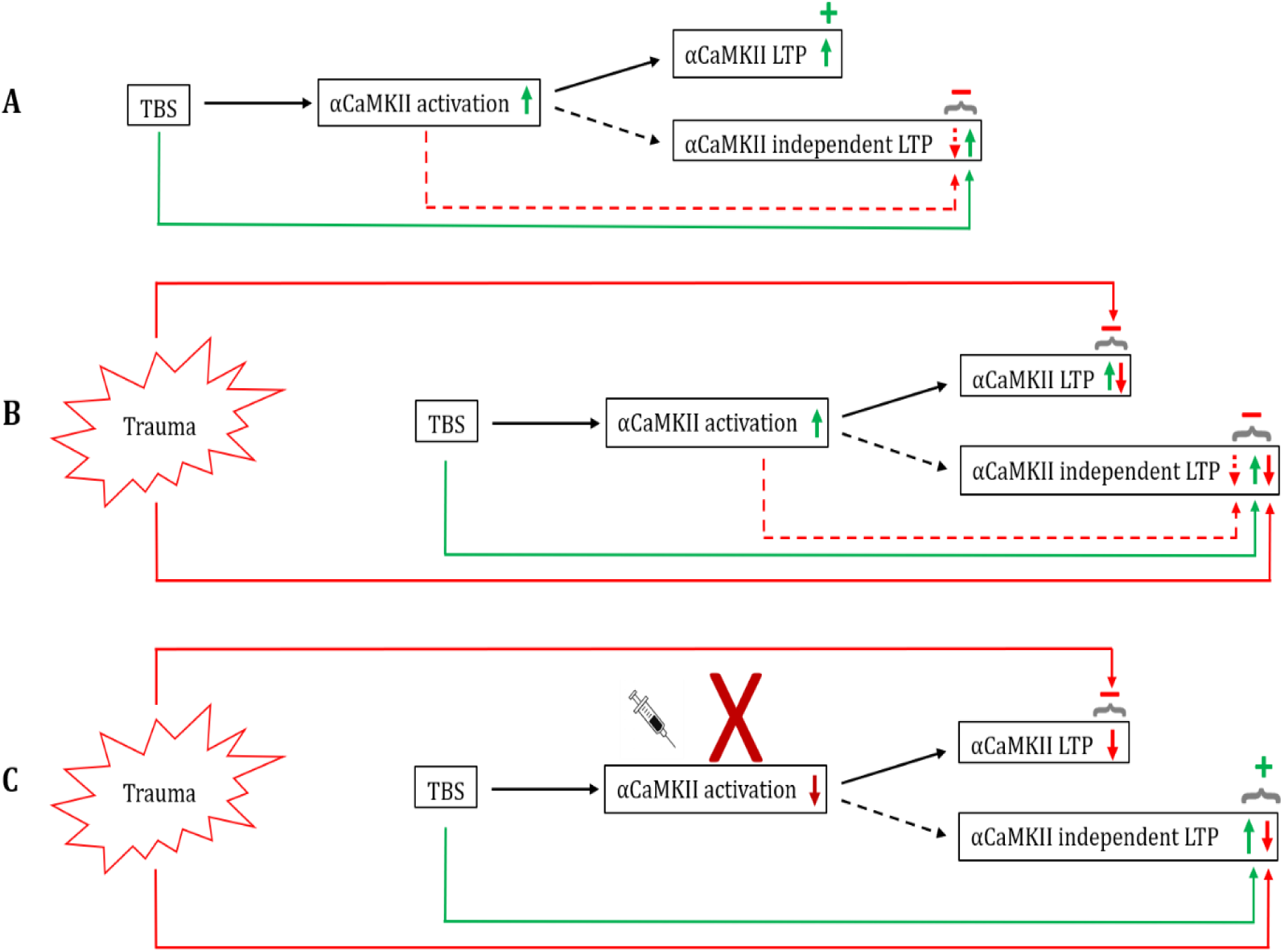
A schematic model illustrating the versatile role of αCaMKII in trauma induced hippocampal plasticity. (A) Under normal conditions, the activation of αCAMKII has a dual effect supporting the formation of αCAMKII-dependent LTP while inhibiting other forms of LTP. As a result, TBS leads mainly to the formation of αCAMKII-dependent LTP. (B) Exposure to trauma tends to suppress both αCAMKII- dependent LTP and αCAMKII independent LTP. (C) Reducing the expression of αCAMKII reduces the probability of activating αCAMKII-dependent LTP. At the same time, it may also remove αCAMKII-induced inhibition of other forms of LTP. Accordingly, it increases the likelihood of inducing these other forms of LTP, despite the inhibitory effect of the trauma exposure.

In summary, the current findings reveal that reducing the level of expression of αCAMKII specifically in the dDG leads to increased resilience in the face of a traumatic experience. These results further emphasize the pivotal role dDG has in stress resilience (27, 56). More generally, the results lend further support to the unique role the dDG has in mediating the interface between emotional and cognitive functions of the hippocampus (8, 24). Accordingly, directed local interventions within the dDG may have the potential to help reducing PTSD symptoms). The current findings also emphasize the contribution of neural plasticity to the normal functioning of the DG and the hippocampus, and the possible contribution of abnormal hippocampal plasticity to trauma-related pathological symptoms. In that respect, the results suggest a relatively novel angle of examining hippocampal plasticity, i.e. the role of healthy or pathological competition between different forms of plasticity in trauma-related pathology and in stress resilience.

## 4. MATERIALS AND METHODS

### 4.1. Animals

Male Sprague-Dawley rats weighing 30-50 gm (Envigo, Jerusalem, Israel) were grouped together (four per cage) upon arrival (postnatal day (PND) 22) and habituated for five days (22 ± 2 °C; light-dark cycle: 12/12 h) with regular food pellets and water ad libitum before commencing the experiments. All experiments were approved by the University of Haifa Animal Care and Use Committee, and performed according to the NIH Guidelines.

### 4.2. Stress Protocols

#### 4.2.1. Juvenile stress (JS)

The juvenile stress protocol was implemented, as previously described (55)(See Supporting Methods for details).

#### 4.2.2. Adulthood stress - Underwater trauma (UWT)

The UWT stress protocol was followed as previously described (55)(See Supporting Methods).

### 4.3. Behavioral assessments

Two weeks after the UWT, rats were behaviorally assessed in the open field (OF) and the elevated plus-maze (EPM). Animal’s behavior was recorded and then analysed using the Etho-Vision XT8 video tracking system (Noldus, Wageningen, Netherlands)

#### 4.3.1. Trauma reminder

Odor re-exposure was employed as a reminder cue (55). Before each behavioral test, animals were re-exposed for 30 s to the same vanilla odor used prior to the UWT exposure. Rats were subjected to the behavioral tests immediately after.

#### 4.3.2. Open field (OF) test

Locomotor activity and anxiety-like behavior were tested at PND 77 (27, 55). Anxiety index was measured as the percentage of exploration (distance/ time) in the center divided by total exploration (distance/ time).

#### 4.3.3. Elevated plus maze test (EPM)

Anxiety-like behavior was assessed in the EPM as previously described (27, 55). EPM was carried out 24 hours after the OF test (PND 78). Anxiety index in the EPM was calculated as the percentage of exploration (distance/ time) in the open arm divided by the total exploration of open and closed arms (distance/ time).

#### 4.3.4. Behavioral profiling

To dissociate between trauma “affected” and “unaffected subjects,” we implemented the “behavioral profiling” approach, which refers to the performance of a control, non-exposed group as characterizing normal behavior, and assessed deviation from the NORM (55). Classification criteria were defined according to the control group variability in performance for each behavioral parameter. Based on validated previous experience (27,52,55,56)), we here developed an algorithm (Fig S2), which screened fifteen different behavioral variables from the OF and EPM, which represent activity and anxiety-like behaviors (Table S1). (For a detailed description See Supporting Methods).

### 4.4. Stereotaxic surgery, and Cannula implantation

Stereotaxic surgery) PND 44-45((27) was performed bilaterally in the dDG relative to bregma (AP= - 3.7; Lateral= ± 2.2; Ventral= - 3.6 mm) according to Paxinos and Watson (86). Cannula were placed during the surgery procedure (For details, see Supporting methods).

#### 4.4.1. KN 93 drug microinjection

On PND 63, following habituation, KN-93, a well-known pharmacological inhibitor of CaMKII, was injected locally in the dDG (5.0 μg/ 0.5 μl saline per hemisphere), 15 minutes before subjecting rats to UWT. (87, 88) (For details, see Supporting methods).

### 4.5. Lentivirus production and validation

The design and construction of pAAV-CMV- αCaMK2-P2A-EGFP and pAAV-U6- shRNA-CMV-EGFP vectors were based on (89). Three shRNA sequences were designed and knockdown efficiency of all three was tested in 293 cells. Since naturally there is no αCAMKII expression in 293 cells, both pAAV-CMV-αCaMK2-P2A-EGFP and pAAV-U6-shRNA-CMV-EGFP vectors were transfected in 293 cells. The reduction of αCaMKII in vitro was validated and quantified by western blotting. Furthermore, in vivo validation was performed by western blot analysis following αCaMKII knockdown in the rat dDG (Fig 2 A-C).

#### 4.5.1. Viral injection

On PND 44-45 rats received bilateral microinjections of either AAV vectors: pAAV- U6-shRNA-CMV-EGFP (αCaMKII V) in the dDG to achieve αCaMKII knockdown (kd) or a control virus. Following lowering the syringe, and after 5 min of rest in the target area, 1 µl of viral vector suspension was injected in each hemisphere (0.15 µl/min) through a 10 µl Hamilton syringe (30G) connected to a motorized nanoinjector (Stereotaxic Injector, Stoelting, Wood Dale, USA). The needle remained in the place for ten minutes before being slowly withdrawn.

### 4.6. In vivo electrophysiology and induction of long-term potentiation (LTP)

The electrophysiology procedure was conducted as described before (28). Two TBS stimulation protocols were used - a regular TBS protocol and a strong TBS protocol. *Regular protocol*: a single set of 10 trains, each of 10 pulses, administrated at 100 Hz with 200 ms inter-train interval. *Strong protocol*: three sets of 10 trains, each of 10 pulses, administrated at 100 Hz with 200 ms inter-train interval and 1 min inter-set interval. LTP was recorded for 60 min after TBS, and measured as the difference in EPSP slope before and 60 min after TBS (For details, see Supporting methods).

### 4.7. Brain removal and histology

(For details, see Supporting methods)

### 4.8. Western blot assay

(For details, see Supporting methods)

### 4.9. Experimental groups and design

(For details, see Supporting methods)

### 4.10. Statistical Analysis

Data were analyzed using the IBM SPSS (21) Statistics software (IBM, Armonk, NY, USA). All behavioral, electrophysiological, and immunoblot results were analyzed using independent sample t-test, one way ANOVA, and repeated measures ANOVA followed by post-hoc Bonferroni, as appropriate. For behavioral profiling, Pearson’s chi-square test was used. Appropriate Greenhouse-Geisser or Huynh-Feldt corrections for sphericity issues were applied when necessary. All results are presented as mean ± SEM.

## Acknowledgments

We thank Dr. Rachel Anunu for her technical support, Dr. Silvia Mandel for editing the manuscript and Anaam Kraiyim for her assistance.

