## Supplement table, method and figure for "The role of hippocampal CAMKII in resilience to trauma-related psychopathology"

**FIGURES**

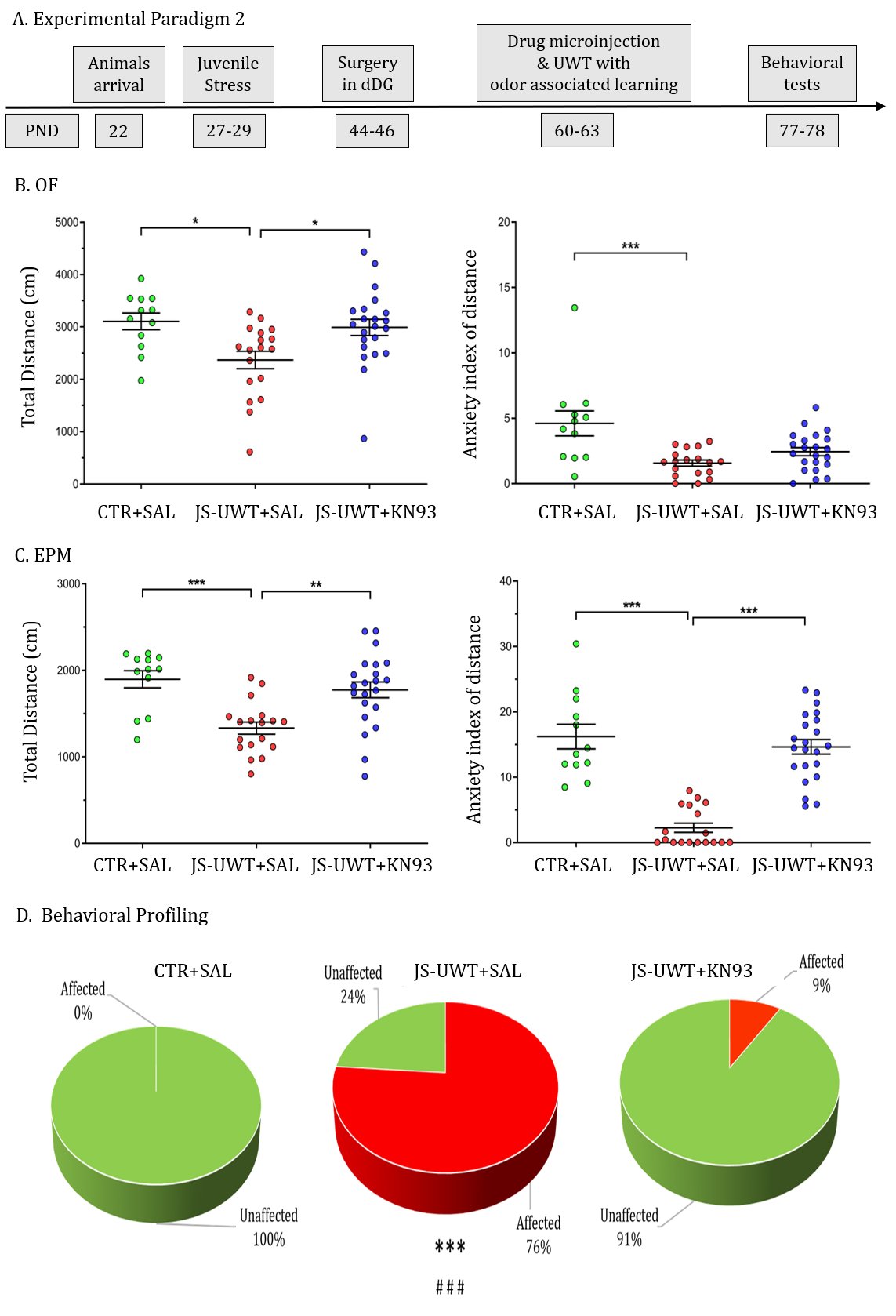

**Figure 1**. **Administration of CaMKII inhibitor KN93 in the dDG reduces the prevalence of affected animals after exposure to juvenile stress and UWT as adulthood stress (JS-UWT).** See supporting information for further details. (A) Schematic representation of experimental timeline, (PND-postnatal day; supporting information). (B) and (C) Averaged group effects in the open field and the elevated plus maze respectively. (Left) Total distance travelled in the maze and (Right) Anxiety index of distance. All values are mean ± SEM ***p<0.001, ** p<0.01, *p<0.05. (D) Behavior profiling analysis shows a significant higher proportion of affected animals among JS-UWT+SAL rats compared to that of the CTR+SAL population, whereas the JS-UWT+KN93 group shows significant reversal effect. Values are the % affected and unaffected animals in each group. ***p<0.001, ###p<0.001.

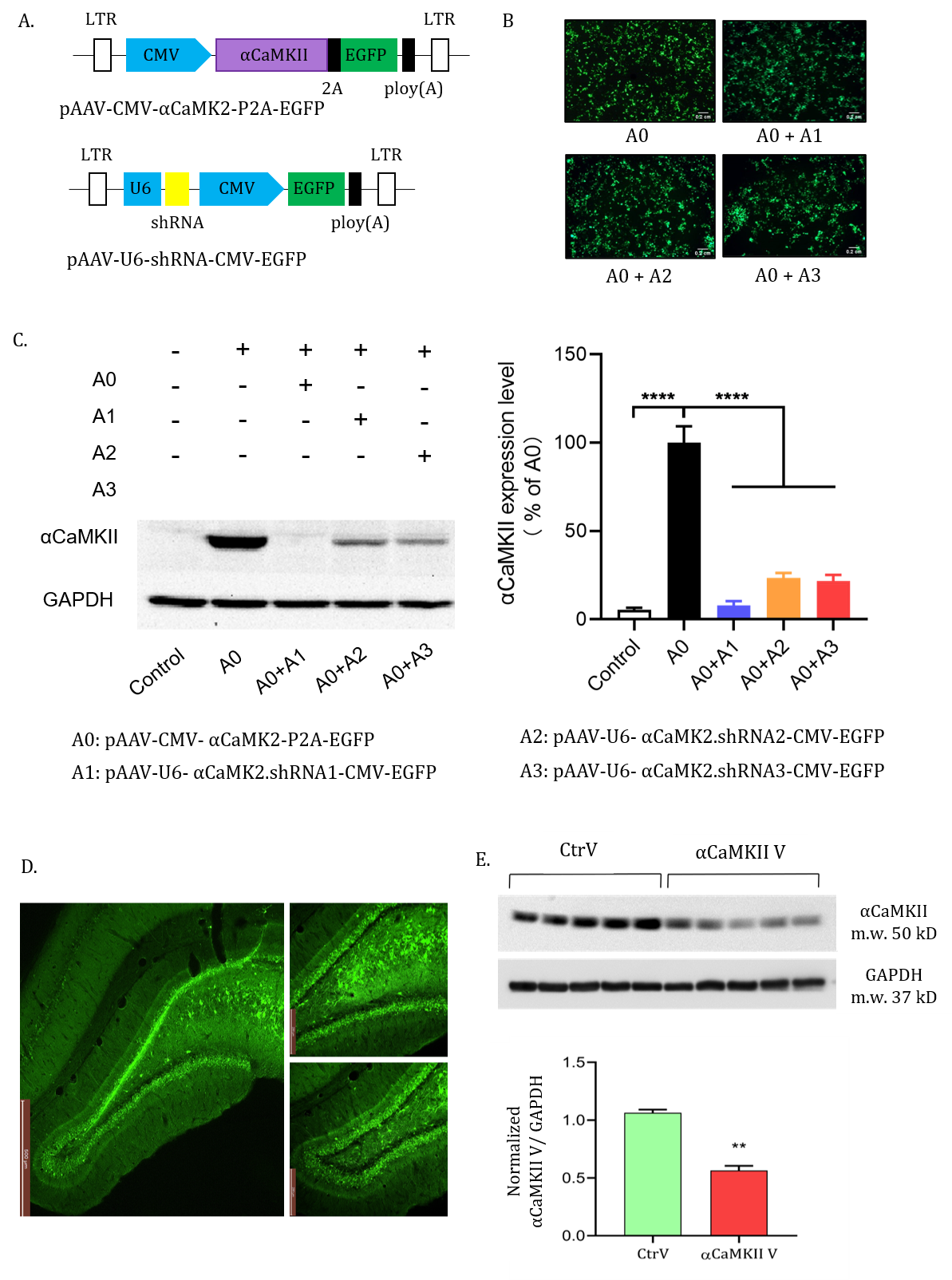

**Figure 2**. **Validation of αCaMKII virus**

The knockdown efficiency of three shRNAs in 293 cell line. (A) up: Schematics of AAV construct for overexpressing (pAAV-CMV-αCaMK2-P2A-EGFP); down: Schematics of AAV construct for knockdown αCaMKII (pAAV-U6-shRNA-CMV-EGFP). ITR, inverted terminal repeats; CMV, cytomegalovirus promoter. (B) Representative fluorescence images of 293 cells after AAV vectors transfection. EGFP, green. (C) αCaMKII protein level in 293 cells transfected with pAAV-CMV-αCaMK2-P2A-EGFP and pAAV-U6-shRNA-CMV-EGFP vectors. (D) αCaMKII virus expression in the dDG. (E) Representative western blot images showed significant reduction in αCaMKII knockdown group. All values are mean ± SEM, ***p<0.001, ** p<0.01, *p<0.05.

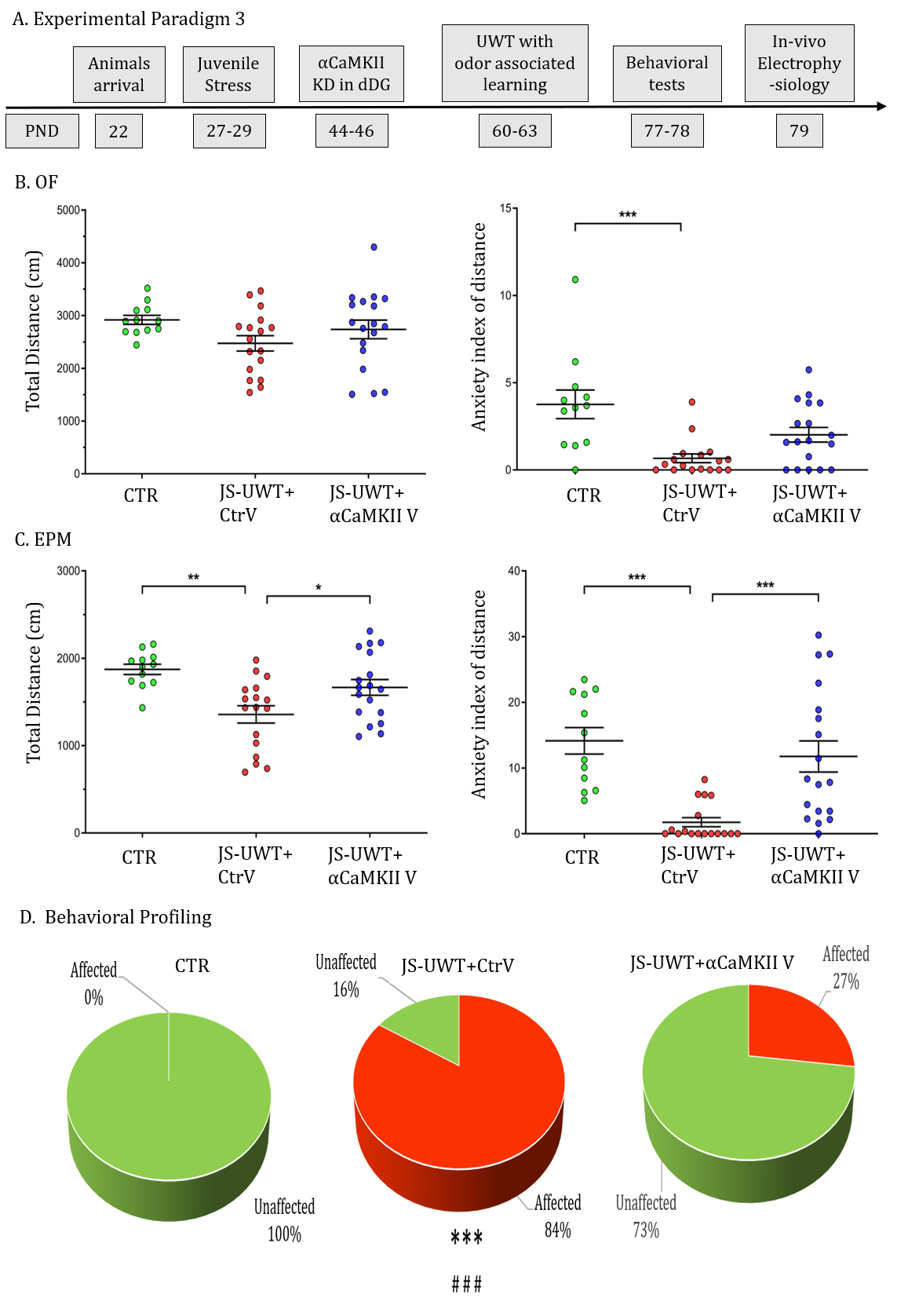

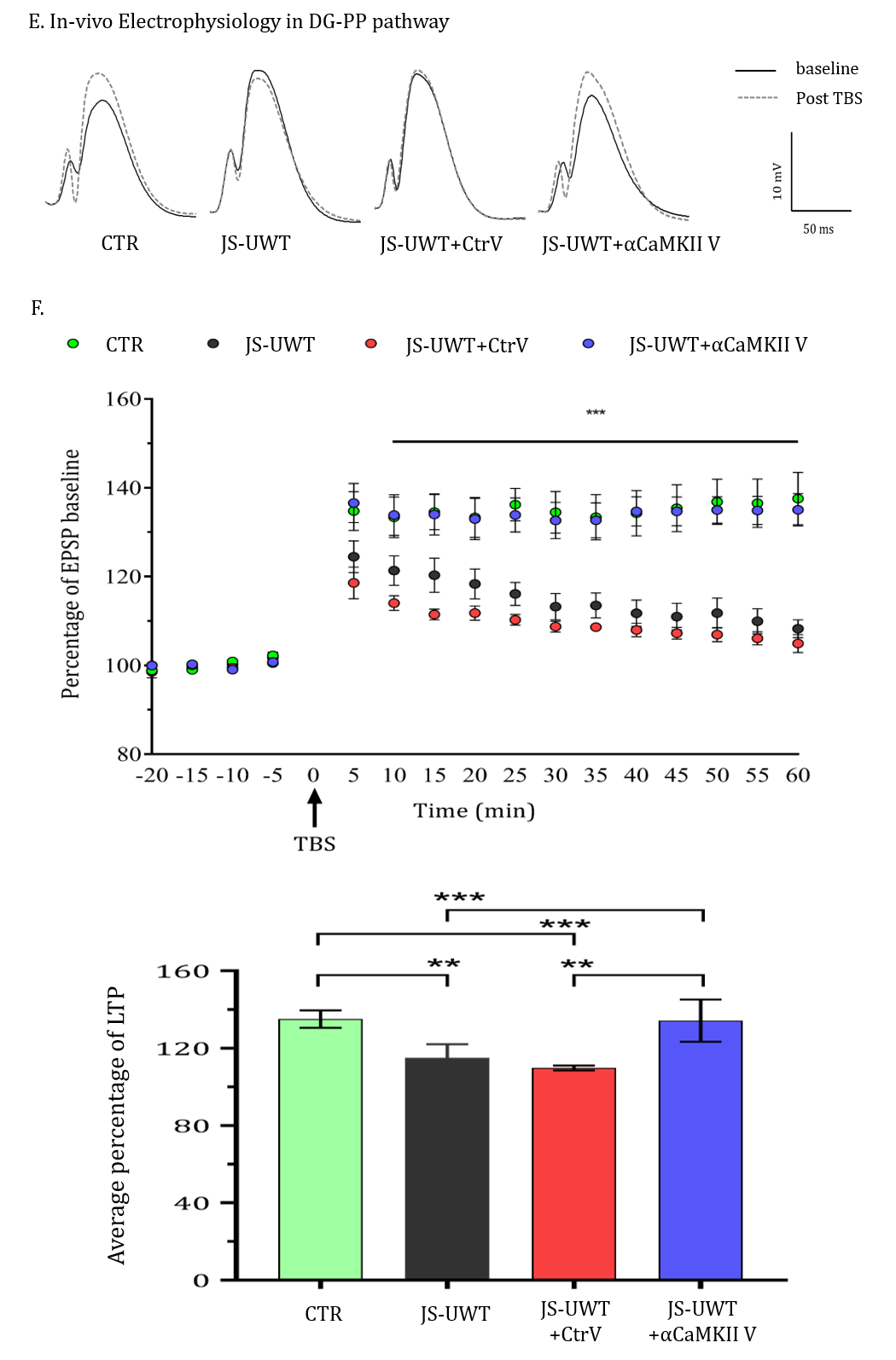

**Figure 3**. **Knock-down of αCaMKII in the dDG reduces the prevalence of affected animals after exposure to JS-UWT and suppressed LTP.** See supporting information for details. (A) Schematic representation of experimental timeline, (PND-postnatal day; supporting information). (B) and (C) Averaged group effects in the open field and the elevated plus maze respectively. (Left) Total distance travelled in the maze and (Right) Anxiety index of distance. All values are mean ± SEM, ***p<0.001, ** p<0.01, *p<0.05. (D) Behavior profile approach shows a significant higher proportion of affected animals among JS-UWT+CtrV rats compared to the CTR population, whereas in JS-UWT+ αCaMKII V group the affected population is significantly less compared to JS-UWT+CtrV group. Values are the % affected and unaffected animals in each group. ***p<0.001, ###p<0.001. (E) Effect of αCaMKII knockdown on synaptic plasticity in the DG of the hippocampus. Evoked field potential response recorded in the DG of CTR (n=8), JS-UWT (n=8), JS-UWT+CtrV (n=10), and JS-UWT+αCaMKII V (n=10) (F) A significant DG field potentiation was recorded after TBS in CTR group and JS-UWT+ αCaMKII V, while TBS failed to induce potentiation in JS-UWT and JS-UWT+CtrV group (left), changes in average percentage of LTP after TBS (right). LTP was assessed upon theta-burst stimulation (TBS) to the perforant path-DG pathway, and recording in the dDG. No significant difference in baseline recording was found prior to TBS. Average percentage of LTP (right), The results are the Mean ± SEM. ***p<0.001, ** p<0.01.

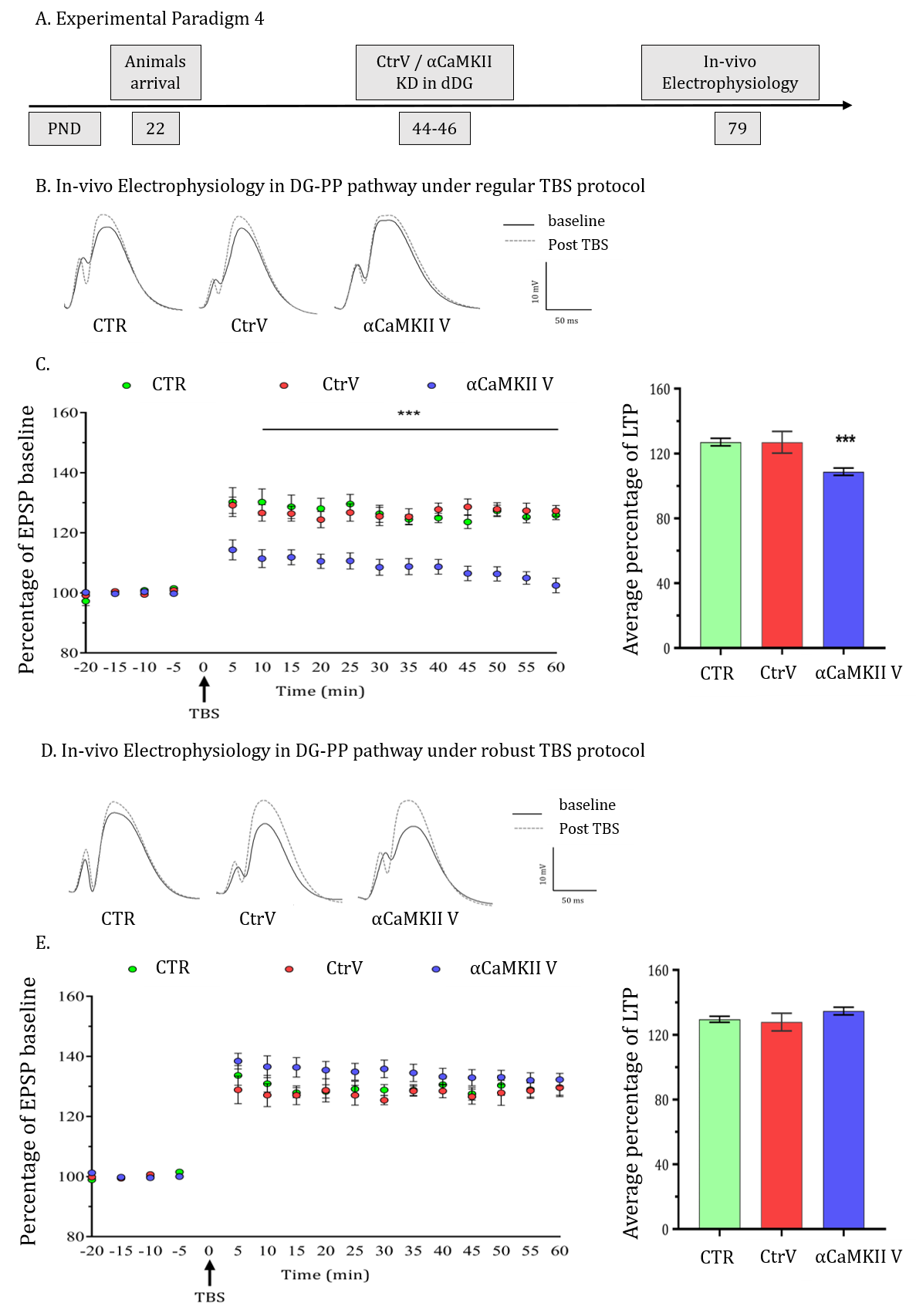

**Figure 4**. **LTP activity under regular or robust TBS protocol following viral inhibition of αCaMKII.** (A) Schematic representation of experimental timeline, (PND-postnatal day; supporting information). (B) Evoked field potential response recorded in the dDG of CTR (n=7), JS-UWT+CtrV (n=7), and JS-UWT+αCaMKII V (n=8) under regular TBS protocol. (C) A significant DG field potentiation was recorded after TBS in CTR and CtrV groups, while in αCaMKII V group, TBS failed to induce potentiation (left); changes in average percentage of LTP after TBS (right). (D) Evoked field potential response recorded in the DG under robust TBS protocol, CTR (n=7), JS-UWT+CtrV (n=7), and JS-UWT+αCaMKII V (n=8). (E) TBS induced a significant potentiation of DG field potentials in all the groups (left); no significant changes were found in average percentage of LTP after TBS (right). LTP was assessed upon TBS to the perforant path-DG pathway, and recording in the dDG. No significant difference in baseline recording was found prior to TBS. The results are the Mean ± SEM, ***p ≤ 0.001.

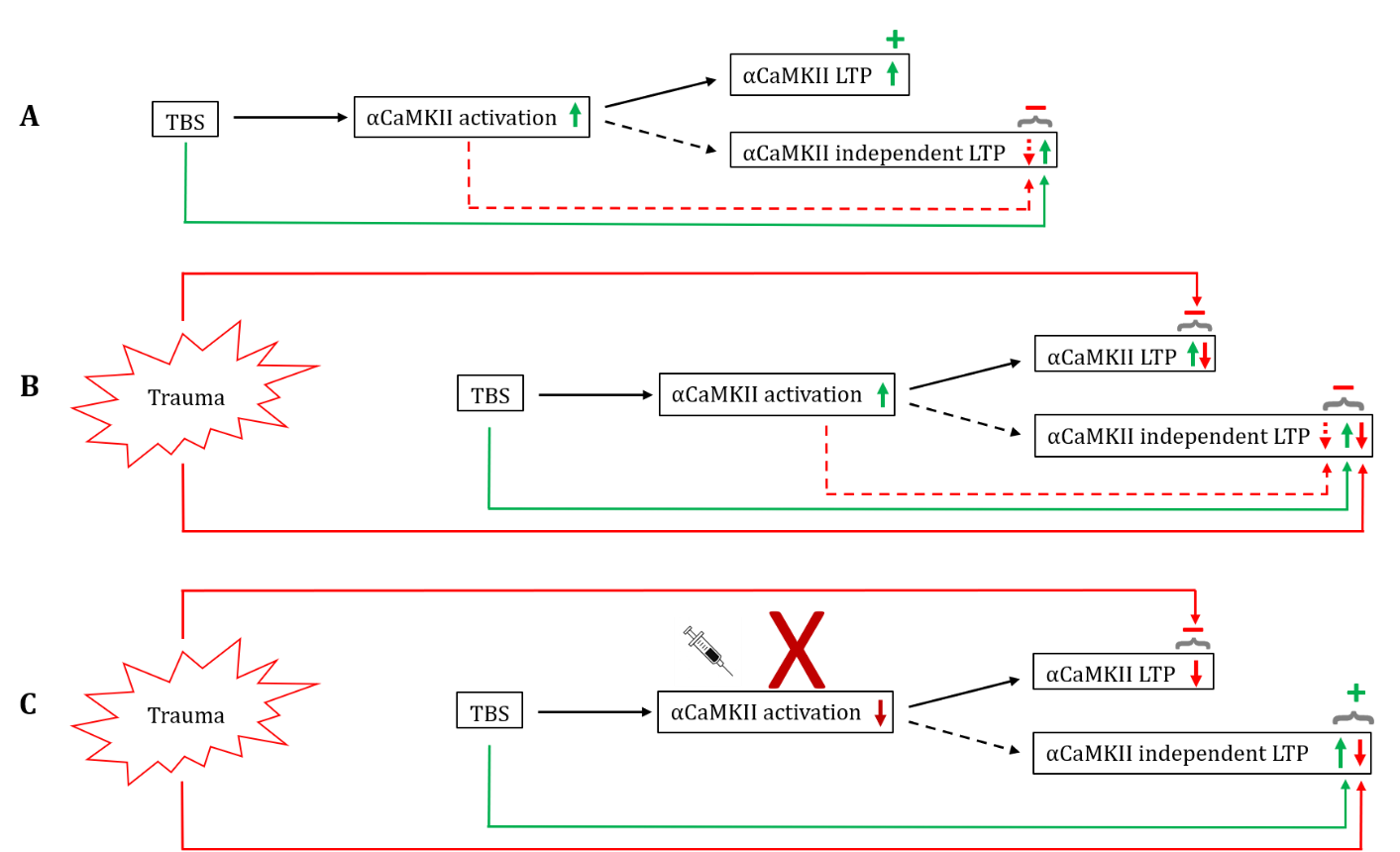

**Figure 5**. **A schematic model illustrating the versatile role of αCaMKII in trauma induced hippocampal plasticity** (A) Under normal conditions, the activation of αCAMKII has a dual effect supporting the formation of αCAMKII-dependent LTP while inhibiting other forms of LTP. As a result, TBS leads mainly to the formation of αCAMKII-dependent LTP. (B) Exposure to trauma tends to suppress both αCAMKII-dependent LTP and αCAMKII independent LTP. (C) Reducing the expression of αCAMKII reduces the probability of activating αCAMKII-dependent LTP. At the same time, it may also remove αCAMKII-induced inhibition of other forms of LTP. Accordingly, it increases the likelihood of inducing these other forms of LTP, despite the inhibitory effect of the trauma exposure.
